# Witnessed trauma exposure induces fear in mice through a reduction in endogenous neurosteroid synthesis

**DOI:** 10.1101/2023.07.04.547710

**Authors:** Aidan Evans-Strong, Najah Walton, Abby Roper, S. Tiffany Donaldson, Mike Lewis, Jamie Maguire

**Author notes:** Corresponding Author: Jamie Maguire, Tufts University School of Medicine Neuroscience Department, 136 Harrison Ave., Boston, MA 02111.

## Abstract

Neurosteroids have been implicated in the pathophysiology of post-traumatic stress disorder (PTSD). Allopregnanolone is reduced in subsets of individuals with PTSD and has been explored as a novel treatment strategy. Both direct trauma exposure and witnessed trauma are risk factors for PTSD; however, the role of neurosteroids in the behavioral outcomes of these unique experiences has not been explored. Here we investigate whether observational fear is associated with a reduced capacity for endogenous neurosteroidogenesis and the relationship with behavioral outcomes. We demonstrate that both mice directly subjected to the threat (foot shocks) and those witnessing the threat have decreased plasma levels of allopregnanolone. The expression of a key enzyme involved in endogenous neurosteroid synthesis, 5α-reductase type 2, is decreased in the basolateral amygdala (BLA), which is a major emotional processing hub implicated in PTSD. We demonstrate that knockdown of 5α-reductase type 2 exaggerates the behavioral expression of fear in response to witnessed trauma, whereas treatment with an exogenous, synthetic neuroactive steroid GABA_A_ receptor PAM with molecular pharmacology similar to allopregnanolone (SGE-516 [tool compound]) decreases the behavioral response to observational fear. These data implicate impaired endogenous neurosteroidogenesis in the pathophysiology of threat exposure, both direct and witnessed. Further, these data suggest that treatment with exogenous 5α-reduced neurosteroids or targeting endogenous neurosteroidogenesis may be beneficial for the treatment of individuals with PTSD, whether resulting from direct or witnessed trauma.

## 1. Introduction

Previous trauma exposure is a major risk factor for psychiatric illnesses, including depression, anxiety and post-traumatic stress disorder (PTSD) (Yehuda et al., 2015). PTSD is characterized by repeated intrusive memories of the trauma, dissociative episodes, and heightened negative reactions to stimuli associated with the trauma, among other symptoms (Association, 2013). Those diagnosed with PTSD are also often afflicted by negative changes in mood and an increase in generalized anxiety. However, not everyone who is exposed to a traumatic event will go on to develop PTSD; current estimates find that 25-30% of those who experience a trauma develop PTSD, putting its prevalence in the population at 7.8% (Gradus, 2017; Jeong et al., 2020; Kessler, Sonnega, Bromet, Hughes, & Nelson, 1995; Yehuda, Resnick, Kahana, & Giller). The factors contributing to the individual differences in the vulnerability to developing PTSD are not yet fully understood.

Both direct trauma exposure and witnessed trauma exposure have been shown to increase risk for PTSD and while the incidence of PTSD following witnessed trauma is lower, the overall prevalence is larger making it a significant societal issue (Yehuda et al., 2015). Various animal models, such as observational fear, have been developed to study the mechanisms through which witnessed trauma increases the risk for psychiatric illnesses (Patki, Solanki, & Salim, 2014; Shi, Feng, Shi, Fu, & Zhou, 2022). Observational fear learning has been employed in rodents and involves pairing a vicarious unconditioned stimulus (US) through observation of another conspecific receiving foot shocks within a specific context. This paradigm has been suggested to model fear transference and affective empathy (Keum & Shin, 2019). These studies have shown that witness trauma in the absence of physical trauma is sufficient to produce long-term behavioral and biochemical changes associated with depression and anxiety in rats and mice (Patki, Salvi, Liu, & Salim, 2015; Warren et al., 2013). Here we employ this paradigm to examine the mechanisms contributing to negative behavioral outcomes following observational fear.

Neuroactive steroids (NAS), which act as positive allosteric modulators of GABA_A_ receptors (GABA_A_Rs), have been implicated in the pathophysiology of PTSD in both clinical and preclinical studies (Almeida, Barros, & Pinna, 2021; Ann M. Rasmusson et al., 2019). Accumulating evidence suggests that altered levels of allopregnanolone may contribute to the pathophysiology of PTSD in at least some subpopulations (Almeida et al., 2021; Pinna, 2019; Schüle et al., 2011). For example, allopregnanolone levels are reduced in the cerebrospinal fluid (CSF) in women with PTSD, which is inversely correlated with PTSD symptoms and negative mood and are most prominent in individuals with comorbid major depressive disorder (MDD) (A. M. Rasmusson et al., 2006). Neuroactive steroids have also been suggested to have therapeutic potential for individuals with PTSD. Unfortunately, treatment with ganaxolone failed to perform better than placebo in a phase 2, proof-of-concept clinical trial which may have been attributable to the lower than the anticipated therapeutic level of ganaxolone (Ann M. Rasmusson et al., 2017). Despite this setback, interest remains in the pathophysiological role of neurosteroids in PTSD.

Endogenous allopregnanolone can be synthesized de novo from cholesterol or through metabolism of the precursor progesterone by the sequential actions of two enzymes, 5α-reductase and 3α-hydroxysteroid dehydrogenase (3α-HSD) (for review see (Silvia Diviccaro, Cioffi, Falvo, Giatti, & Melcangi, 2022)). Two isoforms of 5αLJreductase, type 1 and type 2, are the rate-limiting enzymes involved in neurosteroidogenesis. However, few studies have investigated the potential pathophysiological role of deficits in endogenous neurosteroidogenesis in mood disorders. Inhibition of 5α-reductase with finasteride is associated with long-term deficits in mood, including anxiety and depression, termed post-finasteride syndrome (PFS), which is thought to involve deficits in endogenous neurosteroidogenesis (Silvia Diviccaro, Melcangi, & Giatti, 2020). Further, a polymorphism in the gene encoding for 5α-reductase type 2, SRD5A2, has been associated with an increased risk for PTSD in males (Gillespie et al., 2013). These correlational studies implicate deficits in endogenous neurosteroids in the pathophysiology of PTSD; however, additional studies are required to directly link deficits in endogenous neurosteroidogenesis and PTSD.

Here we investigate whether observational fear is associated with a reduced capacity for endogenous neurosteroidogenesis and its relationship with behavioral outcomes. Observational fear exposure reduces the expression of 5α-reductase type 2 in the basolateral amygdala (BLA) and lowers allopregnanolone levels in both mice that witnessed and were directly subjected to the threat. Mice with impaired endogenous neurosteroidogenesis, either pharmacological or genetic, display an exaggerated fear response to the witnessed trauma, implicating these changes in the observational fear response. Treatment with a synthetic neuroactive steroid GABA_A_R PAM with molecular pharmacology similar to allopregnanolone (SGE-516 [tool compound]) decreased the behavioral response to observational fear. These data demonstrate a role for endogenous 5α-reduced neurosteroidogenesis in the pathophysiology of observational fear and suggest that targeting this pathway may be beneficial for PTSD treatment.

## 2. Material and methods

### 2.1 Animals

Adult (12 weeks old) C57BL/6J mice (Jackson Labs 000664) were used for these studies. Animals were housed on a 12:12 h light/dark cycle at Tufts University School of Medicine in a temperature and humidityLJcontrolled environment with ad libitum access to food and water. All animal procedures were handled according to the protocols approved by the Tufts University Institutional Animal Care and Use Committee (IACUC). Floxed Srd5a2 mice were generated by GenOway (Supplemental Figure 1).

These mice were genotyped using the following primers: 5’-AACTTGTAAATCTTTGCCACTCTGCGG-3’ (F), 5’-AGCAGGAGAAATGTAGGTGAGCAGGG-3’ (R).

### 2.2 Observational fear paradigm

Animals were randomly assigned to be either naïve, observers, or demonstrators. Naïve mice were handled by the experimenters similarly to the observers and demonstrators but not exposed to threat exposure (either witnessed or direct). The observational fear paradigm used, described in Fig 1a, consists of three shock sessions ∼24 hours apart, a 24-hour recall timepoint, and a 96-hour recall timepoint. For all shock sessions, the observer and demonstrator mice were placed on separate sides of a fear conditioning box separated by a clear plastic barrier. Following a 30-second habituation period, 2s foot shocks of 0.7 mA were delivered every 28s to the demonstrator mouse only, lasting for a total of 6 minutes. This was repeated with each observer-demonstrator pair once per day over three consecutive days. During recall timepoints, each observer mouse was placed back into the same half of the fear conditioning box without the demonstrator present. For all shock and recall sessions, freezing behavior was recorded and quantified using Ethovision software.

**Figure 1:**
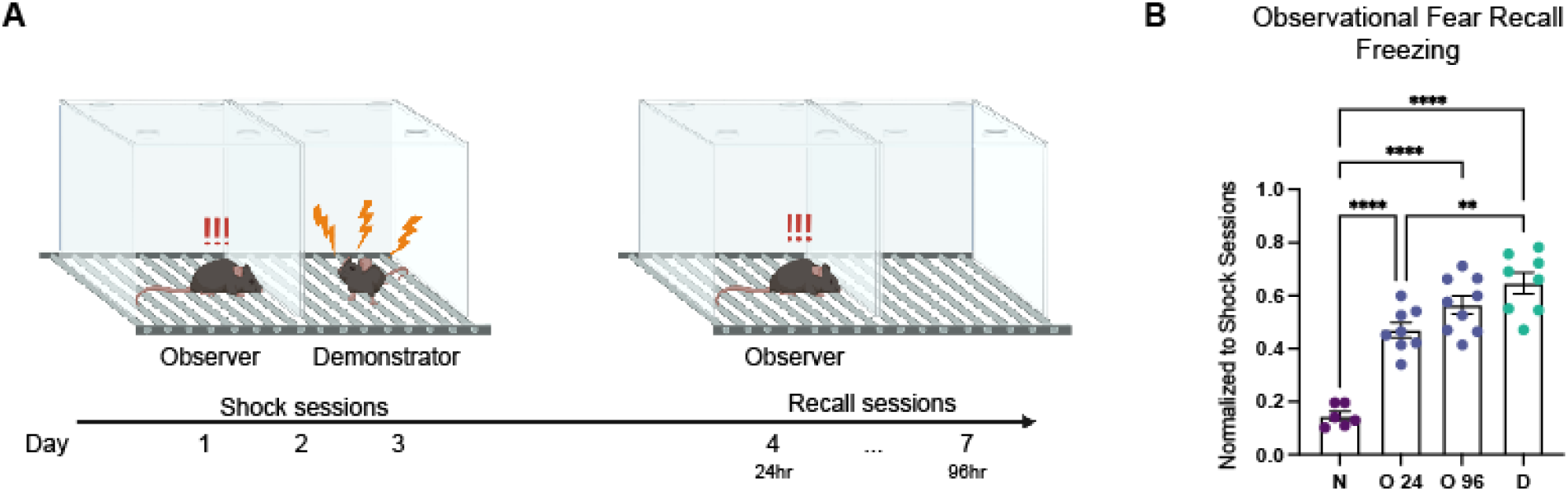
Observational threat exposure produces a contextual fear response. **A** Diagram and schedule of the observational fear paradigm. **B** Normalized freezing times of naïve (N), observer (O) at the 24- and 96-hour timepoints, and demonstrator (D) mice at the 24-hour timepoint. n = 6 – 9 mice per experimental group. [F(3,27)=37.47, R2=0.8063]. ** denotes p<0.01 and **** denotes p<0.0001 using a one-way ANOVA followed by Tukey’s posthoc test.

### 2.3 Drug administration

Finasteride (Sigma-Aldrich F1293) was dissolved in vehicle (15% (2-Hydroxypropyl)-β-cyclodextrin (Sigma-Aldrich H107) in ddH2O) to make a 5 mg/mL solution. Mice were then dosed intraperitoneally at 5 mg/kg according to body weight 24 hours prior to the first shock session and the 24-hour recall session. SGE-516 (450mg/kg/chow) was administered chronically via chow beginning one week prior to the first shock session.

### 2.4 qRT-PCR

Brains were extracted immediately after the final observational fear recall timepoint, flash-frozen in liquid nitrogen, and stored at −80C until sectioning. Tissue punches of the BLA were collected with a 0.5 mm sterile biopsy needle on a cryostat at −20C according to anterior-posterior BLA coordinates from the Allen Mouse Brain Atlas. DNA was extracted and RNA samples were prepared using the SuperScript™ III Platinum™ SYBR™ Green One-Step qRT-PCR Kit (Invitrogen 11736059) as previously described by our laboratory (Darnieder et al., 2019; Walton et al., 2023). Two replicates per animal were measured using primers against 5α2 (sequences: 5’-ATTTGTGTGGCAGAGAGAGG-3’ (F), 5’-TTGATTGACTGCCTGGATGG-3’ (R)), GABA_A_-δ (sequences: 5’-AGGAACGCCATCGTCCTTTT-3’ (F), 5’-CTTGACGACGGGAGATAGCC-3’ (R)), and normalized to β-actin (sequences: 5’-GGCTGTATTCCCCTCCATCG-3’ (F), 5’-CCAGTTGGTAACAATGCCATGTC-3’ (R)) according to previous work from our laboratory. Samples were run using the Mx3000P qPCR system and analyzed with the MxPro software.

Results across duplicates were averaged, and values of experimental genes were normalized to β-tubulin values for each sample. Fold-change from naïve mice was then calculated to determine the changes in RNA transcript levels following observational fear in both observers and demonstrators.

### 2.5 Allopregnanolone measurements

For allopregnanolone measurements, serum was collected immediately after observational fear exposure and allopregnanolone levels were measured by liquid chromatography-tandem mass spectrometry (LC-MS/MS) by PharmaCadence as previously described (Walton et al., 2023). Allopregnanolone levels in the range of 0.2 to 200 ng/ml were measured against reference material purchased (Tocris Cat No. 3653) and a penta-deuterated stable isotope labeled internal standard (Tocris Cat No. 5532). Samples were prepared using liquid-liquid extraction with chlorobutane and diluted 1:1 with deionized water prior to extraction with 1 mL of chlorobutane. The organic layer was extracted, dried under a stream of nitrogen, reconstituted in a 1:1 mixture of deionized water and acetonitrile, and detection was accomplished using a Sciex 5500 Qtrap mass spectrometer operated in positive ion mode. Concentrations were calculated using the integrated peak area and the response factor determined by 1/concentration² weighted, least-squares regression of the calibration curve.

### 2.6 Statistical tests

Statistical analysis was performed using GraphPad Prism 9 software. Unpaired Student’s t-tests were used to compare two experimental groups and one-way ANOVAs were used to compare more than two experimental groups. Tukey’s post-hoc test employed to analyze individual group differences. One-sample t-tests were used to analyze fold change values. Values in this paper are written as mean ± SEM. P-values of < than 0.05 were considered statistically significant and denoted as p ≥0.05=n.s., p<0.05=*, p<0.01=**, p<0.001=***, and p<0.0001=****.

## 3. Results

### 3.1 Observational threat exposure produces a contextual fear response

The observational fear (OF) paradigm has been widely used as an animal model of fear transference as well as empathy (Jeong et al., 2020; Keum & Shin, 2019; Pisansky, Hanson, Gottesman, & Gewirtz, 2017; Shi et al., 2022). Our model was adapted from (Jeon & Shin, 2011) and (Davis, Zaki, Maguire, & Reijmers, 2017) and involves three consecutive days of shock sessions in which the observer mouse witnesses a demonstrator mouse undergoing a series of foot shocks. The animals are separated by a clear plastic barrier and thus the observer can receive visual, auditory and olfactory cues from the demonstrator throughout the threat exposure. In accordance with previously published data (Shi et al., 2022), we find that observer mice show significantly increased freezing behavior (.47 ± .03 normalized to shock session baselines) when returned to the chamber for contextual recall as compared to naïve mice (.15 ± .02) (p < .0001, Tukey’s test, Fig 1b). Interestingly, the behavioral expression of fear following witnessed trauma is nearly the same magnitude as the animals that directly experienced the foot shocks (demonstrator: .65 ± .04 normalized to shock session baselines) (p = .0035, Tukey’s test, Fig 1b). Thus, this model is valuable for examining the neurobiological processes contributing to the pathophysiological consequences of witnessed trauma exposure.

### 3.2 Observational threat impairs endogenous neurosteroid signaling

In order to investigate the hypothesis that altered endogenous neurosteroids may contribute to the behavioral expression of observation fear, we measured the levels of endogenous allopregnanolone in the plasma from mice subjected to observational fear compared to naïve mice and demonstrators. Plasma levels of allopregnanolone are reduced in both observer (1.138 ± .113 ng/ml) and demonstrator (1.044 ± .157 ng/ml) mice compared to naïve controls (2.035 ± .205 ng/ml) (observers: p = .042, demonstrators: p = .0028, one-way ANOVA, Fig 2a). To determine whether endogenous neurosteroid levels in the brain may also be impaired, we measured the expression levels of two key neurosteroidogenic enzymes: 5α-reductase type 1 and type 2. We found a significant decrease in the transcript expression of Srd5a2 in the BLA of observer mice (.393 ± .099 fold change from naïve) (p = .0005, one sample t-test) compared to naïve (Fig 2b). Interestingly, transcript levels of Srd5a2 in the BLA were reduced to a comparable amount in observer (.393 ± .099) and demonstrator mice (.174 ± .049) (p = .209, one way ANOVA). However, we found no significant change in the levels of Srd5a1 transcript in the BLA in observer (.709 ± .297 fold change from naïve) or demonstrator (.843 ± .166) mice (p = .911, one way ANOVA and one sample t-test, Fig S2) compared to naïve. Allopregnanolone has been shown to exert effects on emotional processing through actions on δ-subunit containing GABA_A_ receptors (GABA_A_Rs) in the BLA (Antonoudiou et al., 2022; Silvia Diviccaro et al., 2022). We found that transcript levels of the GABA-δ receptor (Gabrd) were reduced in the BLA of observer (.288 ± .047 fold change from naïve) as well as demonstrator (.200 ± .022) mice compared to naïve (p < .0001, one sample t-test, Fig 2c). These data suggest that endogenous neurosteroid synthesis and signaling is compromised following observational fear exposure.

**Figure 2:**
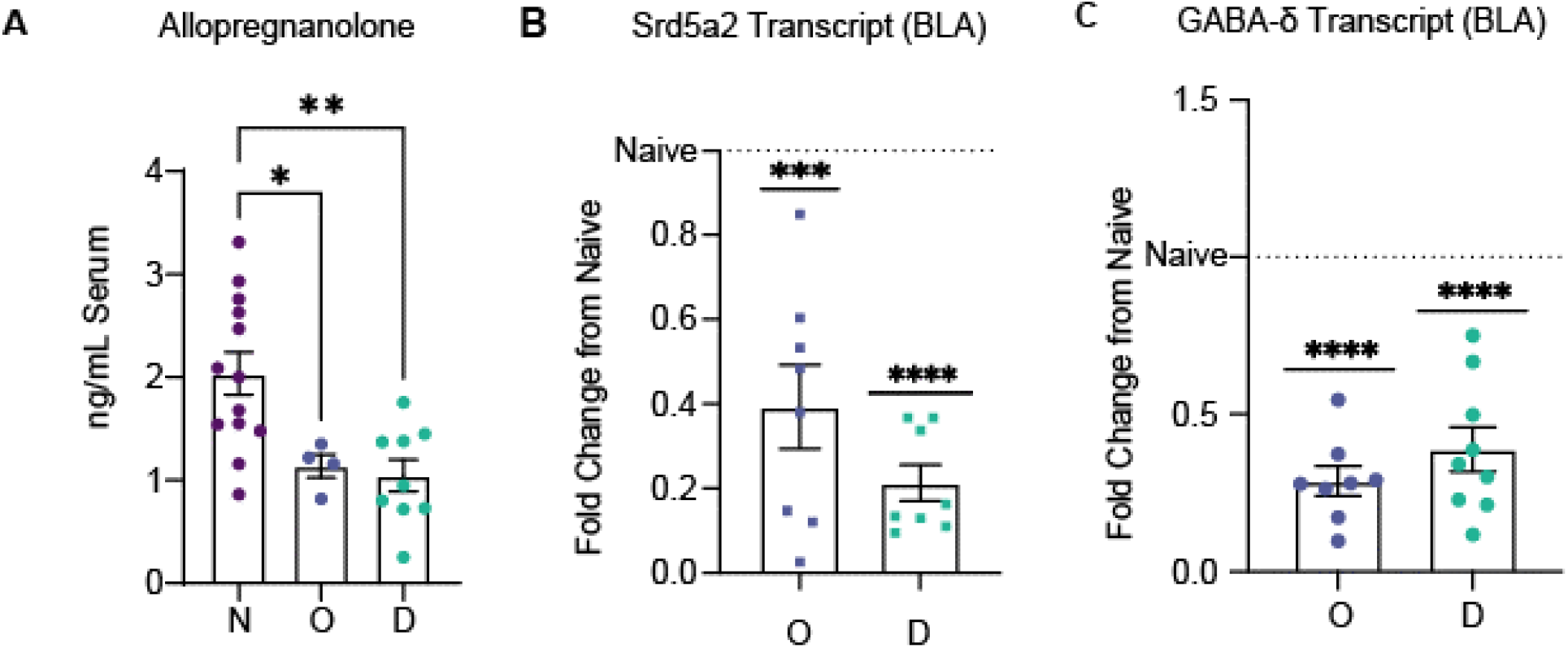
Observational threat impairs endogenous neurosteroid signaling. **A** Serum levels of allopregnanolone in observer (O), demonstrator (D) and naïve (N) mice after observational fear exposure. n = 4 – 13 mice per experimental group. [F (2,23)=3.555, R2=0.4156]. p<0.05=* and p<0.01=** using an ordinary one-way ANOVA followed by Tukey’s posthoc test **B** Transcript levels of Srd5a2 in the BLA in observer (O) and demonstrator (D) mice after undergoing observational fear. Transcript levels are shown normalized to naïve average (1.0).**C** Transcript levels of Gabrd in the BLA in observer (O) and demonstrator (D) mice after undergoing observational fear. Transcript levels normalized to naive average (1.0).p<0.001=*** and p<0.0001=**** using a one-sample t test.

### 3.3 Impaired endogenous neurosteroidogenesis alters the contextual observational fear response

To directly investigate the role of endogenous neurosteroid signaling on the behavioral expression of observational fear, we generated a novel mouse model which enables us to manipulate 5α-reductase type 2 expression. Based on the impairment of Srd5a2 expression following observational fear exposure, we crossed floxed Srd5a2 mice generated by genOway with CamKII-Cre mice to knockout this key neurosteroidogenic enzyme in excitatory neurons, which was sufficient to reduce Srd5a2 protein levels in the BLA (.343 ± .025 fold change from naïve) compared to controls (Fig 3b; p = .090, ANOVA). Mice lacking Srd5a2 in CamKII neurons (Srd5a2/CamKII) exhibited increased behavioral expression of observed fear, increasing freezing behavior during recall (1.121 ± .146 normalized to shock session baselines) compared to controls (.590 ± .076) (Fig 3c; p = .017, Welch’s T test).

**Figure 3:**
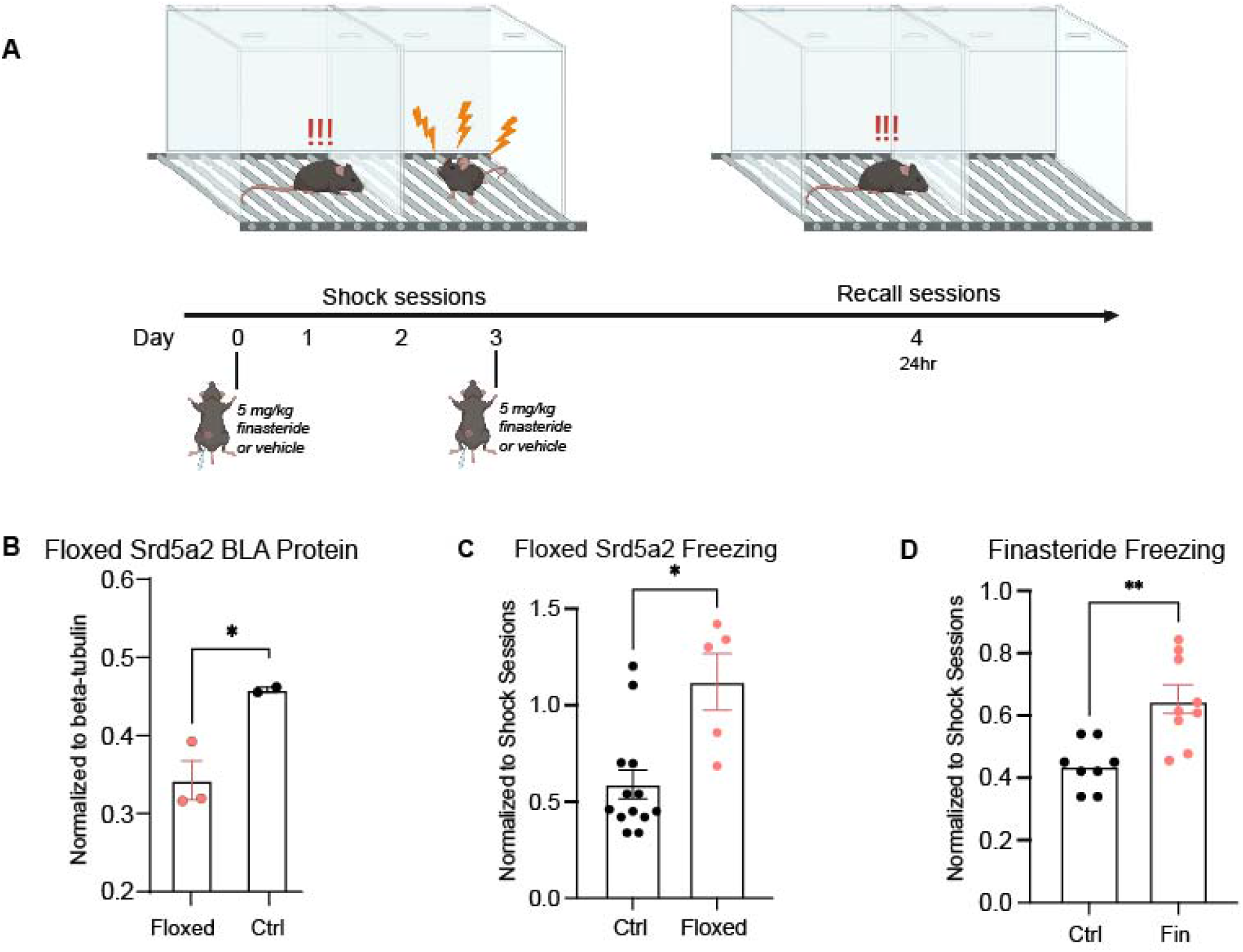
Impaired endogenous neurosteroidogenesis alters the contextual observational fear response. **A** Timeline of the observational fear paradigm including finasteride dosing 24 hours prior to the first shock session and the recall session. **B** Protein expression of 5_α_-reductase type 2 in the BLA of heterozygous floxed (floxed) mice compared to wild type (Cre-) controls (Ctrl). **C** Floxed Srd5a2 mice exhibit an increase in freezing behavior during 24hr recall following observational fear compared to controls. **D** Mice treated with finasteride also exhibit an increase in freezing behavior during 24hr recall compared to controls following observational fear. n = 2-13 mice per experimental group. p<0.05=* and p<0.01=** using a Students t-test.

Finasteride is a pharmacological inhibitor of 5α-reductase, resulting in a reduction in the synthesis of allopregnanolone (Silvia Diviccaro et al., 2020). The impact of finasteride on the behavioral expression of observational fear was measured to further explore the role of endogenous neurosteroid synthesis in the behavioral sequelae of witnessed trauma. Mice treated with finasteride (5 mg/kg) (Kumazaki et al., 2011) exhibited an increase in the behavioral expression of observational fear, increasing their freezing behavior (.646 ± .046 normalized to shock session baselines) compared to vehicle treated mice (.439 ± .027) (Fig 3d; p = .002, Welch’s T test). Thus, both genetic and pharmacological inhibition of 5α-reductase is sufficient to increase the behavioral expression of observational fear, suggesting that the observed reduction in 5α-reductase may contribute to the fear response following observational fear exposure.

### 3.4 Exogenous neurosteroid treatment decreases the observational fear response

Given that our data indicates that observational fear is correlated with decreased allopregnanolone synthesis and/or signaling, it raises the question of whether treatment with a GABA_A_R PAM may reduce the behavioral expression of observational fear. To test this hypothesis, we treated mice with chow containing SGE-516 (450 mg/kg/chow, developed by SAGE Therapeutics) for one week prior to shock sessions (Fig 4a) exhibited decreased freezing at both recall timepoints (.501 ± .049 at 24 hours; .462 ± .047 at 96 hours) as compared to controls (.749 ± .075 at 24 hours; .745 ± .073 at 96 hours) (Fig 4b; p = .010 at 24 hours; p = .0031 at 96 hours; unpaired T tests). These data suggest that NAS GABA_A_R PAMs, such as SGE-516, may have therapeutic utility for witnessed trauma exposure.

**Figure 4:**
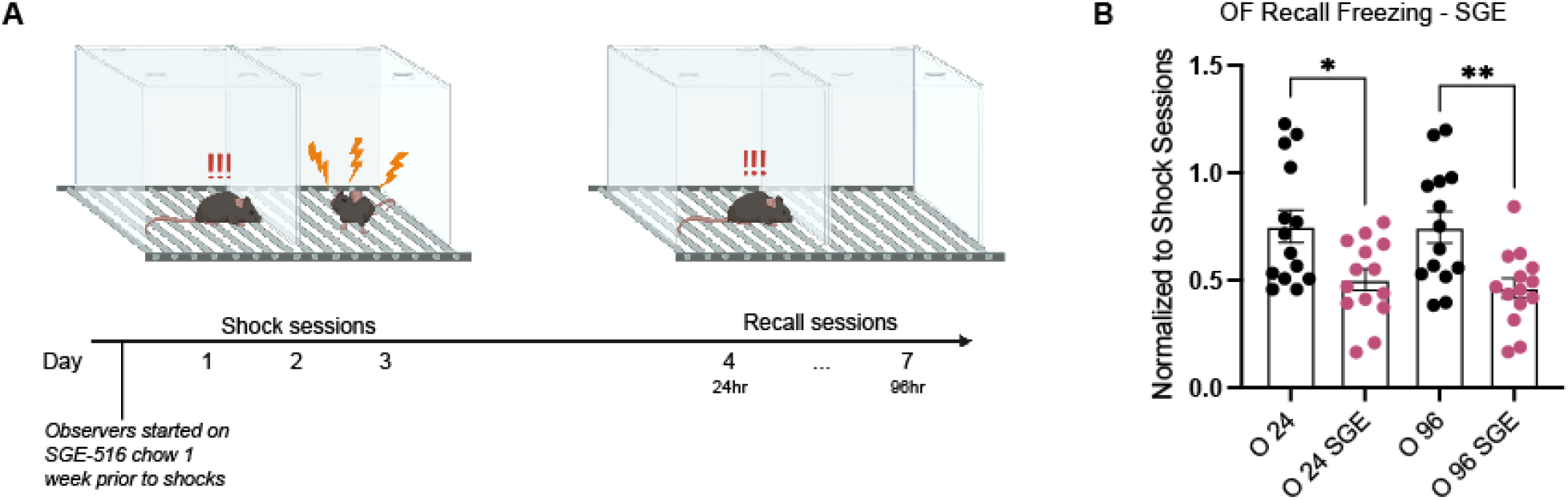
Exogenous neurosteroid treatment decreases the observational fear response. **A** Diagram of the experimental paradigm with schedule of SGE-516 treatment relevant to observational fear exposure. **B** The average freezing behavior of observer mice dosed with vehicle during the 24-and 96-hr recall sessions in vehicle-treated observer mice (O 24 and O 96) and observers treated with SGE-516 (O 24 SGE and O 96 SGE). n = 14 mice per experimental group. p<0.05=*and p<0.01=** using a Students t-test.

## 4. Discussion

### 4.1 Trauma exposure-induced impaired neurosteroid synthesis enhances the behavioral expression of fear

Deficits in allopregnanolone levels have been associated with PTSD (Pinna, 2019) (A. M. Rasmusson et al., 2006). However, these studies were largely correlational and the direct role of endogenous neurosteroids in the pathophysiology of PTSD remains unclear. Here we demonstrate that both witnessed and direct trauma exposure reduces endogenous allopregnanolone levels in mice. These deficits in endogenous allopregnanolone levels are associated with a reduction in the expression of 5α-reductase, a key neurosteroidogenic enzyme, in the BLA. While these changes are still correlational, these alterations are consistent with those observed in individuals with PTSD (Pinna, 2019; Ann M. Rasmusson et al., 2019). In an effort to directly interrogate the role of endogenous neurosteroids on the behavioral sequelae of witnessed trauma, we assessed observational fear responses in mice with a knockdown of 5α-reductase type 2. Mice with a reduced capacity for endogenous neurosteroid synthesis, either genetic or pharmacological, exhibited an exaggerated behavioral expression of fear following observation fear exposure. These data suggest that deficits in endogenous neurosteroids may directly contribute to the behavioral deficits associated with witnessed trauma. Additionally, we demonstrate that treatment with a synthetic NAS GABA_A_R PAM is sufficient to reduce the behavioral expression of fear, further implicating neurosteroid deficits in the pathophysiology of witnessed trauma. The ability of NAS GABA_A_R PAMs to improve behavioral outcomes following observational fear may be due to the ability to modulate BLA network states (Antonoudiou, 2022) which have been demonstrated to govern the behavioral expression of fear (Davis, 2017; Ozawa, 2020). Taken together, these findings indicate that the observational fear response is at least partially driven by inhibition of neurosteroidogenesis in the BLA, and that targeting this mechanism may have therapeutic benefit.

### 4.2 Chronic stress and threat exposure impair endogenous neurosteroidogenesis

The impact of stress on neurosteroids is related to the magnitude and the duration of the stressor. Acute stress has been shown to increase circulating levels of allopregnanolone (Purdy, Morrow, Moore, & Paul, 1991) (Barbaccia et al., 1996) (Barbaccia et al., 1997), which involves at least in part endogenous neurosteroid synthesis in the brain (Purdy et al., 1991). Although few studies have explored the impact of chronic stress on endogenous neurosteroid levels, it is thought that chronic stress and deficits in endogenous neurosteroids contribute to psychiatric illnesses, such as anxiety and depression (for review see (Zorumski, Paul, Izumi, Covey, & Mennerick, 2013) (Almeida et al., 2021)). Protracted social isolation (Agís-Balboa et al., 2007; Dong et al., 2001) and chronic unpredictable stress (Walton et al., 2023) have both been shown to decrease allopregnanolone levels, demonstrating that chronic stress alters endogenous neurosteroid levels. Chronic stress-induced impairment in endogenous neurosteoridogenesis is also supported by the evidence that social isolation and chronic unpredictable stress decrease the expression of 5α-reductase in the brain (Agís-Balboa et al., 2007; Dong et al., 2001; Walton et al., 2023). Impaired endogenous neurosteroid synthesis following chronic stress has been implicated in stress-related behavioral impairments. Consistent with the current findings, the reduction in 5α-reductase expression following social isolation has been associated with an increase in the behavioral expression of fear (Pibiri, Nelson, Guidotti, Costa, & Pinna, 2008).

Previous work from our laboratory has also demonstrated that chronic stress decreases the expression of the GABA_A_R δ subunit, the primary site of action for neurosteroid modulation, in the BLA (Walton et al., 2023). Similarly, chronic stress has been shown to impair tonic GABAergic inhibition in principal neurons in the BLA (Liu et al., 2014). The impact of stress on GABAergic signaling is also related to the extent and magnitude of the stress exposure and is brain region specific (for review see (Maguire, 2014; Maguire & Mody, 2009; Mody & Maguire, 2011)). Related to neurosteroid signaling, acute and chronic stress have been shown to increase GABA_A_R δ subunit and tonic GABAergic inhibition in the hippocampus (Holm et al., 2011; Maguire & Mody, 2007; Matsumoto, Puia, Dong, & Pinna, 2007; Serra et al., 2000; Serra, Pisu, Mostallino, Sanna, & Biggio, 2008) in contrast to the observed decrease in the BLA (Walton et al., 2023). Knockdown of the GABA_A_R δ subunit in the BLA is sufficient to recapitulate the behavioral deficits associated with chronic stress (Walton et al., 2023). Related to the current study, mice which lack the δ subunit of the GABA_A_R exhibit an enhancement of trace fear conditioning (Wiltgen, Sanders, Ferguson, Homanics, & Fanselow, 2005) and allopregnanolone decreases the behavioral expression of contextual fear (Rabinowitz, Cohen, Finn, & Stackman, 2014). These data are consistent with our current findings demonstrating a decrease in 5α-reductase expression following observational fear and that treatment with a NAS GABA_A_R PAM decreases the behavioral expression of witnessed fear.

### 4.3 Endogenous neurosteroids and affective tone

Finasteride is a selective inhibitor of 5α-reductase and was approved for the treatment of lower urinary tract symptoms associated with benign prostatic hyperplasia in 1992 and androgenetic alopecia in early 2000 (for review see (Traish, Melcangi, Bortolato, Garcia-Segura, & Zitzmann, 2015)). The number of annual prescriptions of finasteride in the United States has exceeded over 2.4 million in 2020. This is alarming given the well-documented and serious side effects of finasteride, which includes sexual dysfunction, depression, diabetes, high grade prostate cancer and vascular disease (for review see (Silvia Diviccaro et al., 2020; Traish et al., 2015)). The affective side effects of these compounds are so severe that there was a request to the FDA to pull the drug from the market, which was rejected but now requires a disclosure about the increased suicide risk. The most frequent side effects of finasteride treatment are classified into the following categories: physical, sexual, cognitive and psychological effects with depression and anxiety reported as the most common psychiatric complications (for review see (Alhetheli, Alrashidi, & Alhammad, 2022; Hirshburg, Kelsey, Therrien, Gavino, & Reichenberg, 2016; Traish, 2020)).

Sexual side effects have been reported in 3.4-15.8 percent of men (Hirshburg et al., 2016) which is thought to be a gross underestimate of the actual problem (Giatti, Diviccaro, Panzica, & Melcangi, 2018). Several studies have demonstrated an association between finasteride use and the incidence of depression (Altomare & Capella, 2002; Irwig, 2012; Rahimi-Ardabili, Pourandarjani, Habibollahi, & Mualeki, 2006; Welk et al., 2017) (for review see (Hirshburg et al., 2016)). Alarmingly, former finasteride users had a marked increase in suicidal thoughts (44%) compared to non-users (3%) (Irwig, 2012) which may be particularly relevant for a subset of patients with a history of mood disorders (Laanani, Weill, Jollant, Zureik, & Dray-Spira, 2023). Fewer studies have explored the relationship between anxiety and finasteride usage but self-reports indicate a comparable number of patients experiencing anxiety (73%) as depression (74%) (Walf, Kaurejo, & Frye, 2018). Despite the patient outcry (Walf et al., 2018), accumulating evidence (for review see (Alhetheli et al., 2022; Hirshburg et al., 2016; Traish, 2020)) and high potential cost to benefit of the use of these compounds, there remain skeptics and advocates for their continued use (Rezende, Dias, & Trüeb, 2018) when in fact this is a significant issue for patients and a “surmountable challenge for clinicians” (Traish, 2020).

Consistent with these clinical findings, preclinical studies demonstrate that finasteride treatment alters valence processing. Finasteride treatment increases negative valence processing, increasing stress-induced helplessness in the forced swim test and increasing avoidance behaviors in the open field and elevated plus maze (S. Diviccaro et al., 2019; Godar et al., 2019; Sasibhushana, Shankaranarayana Rao, & Srikumar, 2019). Finasteride treatment also impairs positive valence processing, inducing anhedonia in rats (Godar et al., 2019). Similar to the effects of pharmacological inhibition of 5α-reductase, genetic knockdown of 5α-reductase in the BLA is sufficient to increase negative valence processing, increasing stress-induced helplessness and increasing avoidance behaviors (Walton et al., 2023). Conversely, overexpression of 5α-reductase is sufficient to prevent the behavioral deficits, including increased negative valence processing, associated with chronic stress exposure (Walton et al., 2023). These data provide empirical evidence supporting the clinical associations between 5α-reductase inhibition and psychiatric complications and are consistent with our current findings demonstrating that impaired endogenous neurosteroid synthesis (either genetic or pharmacological) increases negative valence processing. Further, it demonstrates a role for endogenous neurosteroid signaling in setting a baseline affective tone.

### 4.5 Therapeutic potential of harnessing endogenous neurosteroidogenesis

The current study implicates impaired endogenous neurosteroid synthesis in the pathophysiology of witnessed trauma. Previous studies also demonstrated a reduced capacity for endogenous neurosteroid synthesis in the behavioral deficits associated with chronic stress (Walton et al., 2023). Interestingly, these studies also demonstrated that increasing endogenous neurosteroidogenesis was capable of preventing the behavioral deficits following chronic stress (Walton et al., 2023), suggesting that enhancing endogenous neurosteroidogenesis may have therapeutic potential. Treatment with exogenous allopregnanolone has demonstrated clinical antidepressant effects and in the current study is sufficient to reduce the behavioral expression of fear following witnessed trauma. Rather than administering exogenous allopregnanolone, a superior therapeutic approach may be to target 5α-reductase, implicated in the underlying neurobiology, to enhance endogenous neurosteroid synthesis. In addition, 5α-reductase may serve as a useful biomarker for identifying those at risk for developing PTSD as well as those that may be amenable to neurosteroid-based treatments. Further studies are required to explore the translational potential of targeting 5α-reductase for treatment.

---

**Supplemental Figure 1:**
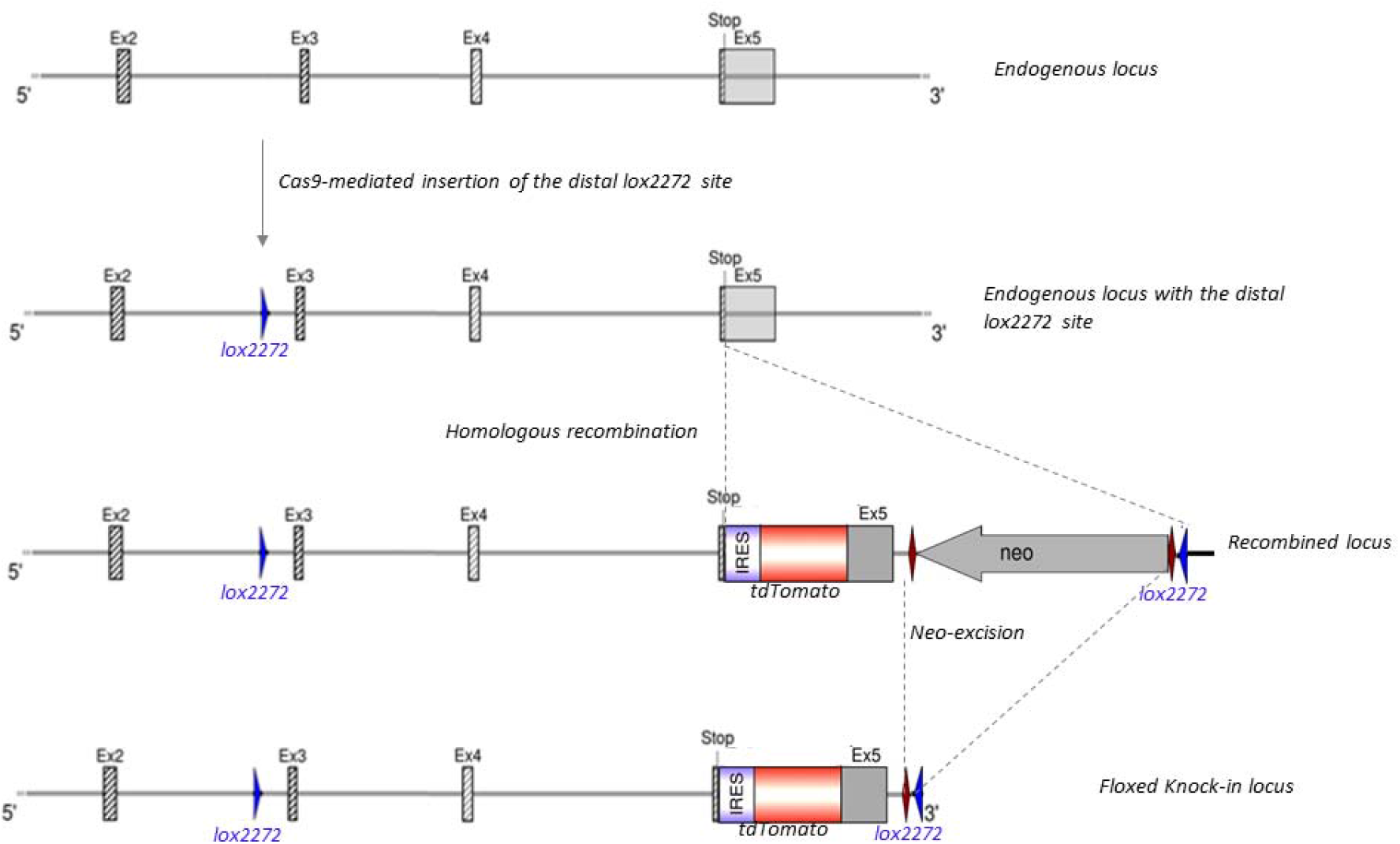
Floxed Srd5a2 mouse model. CRISPR Cas9-mediated homologous recombination was used to introduce a tdTomato reporter and loxp sites between Exon 3 and Exon 5. This approach enables us to visualize 5_α_-reductase type 2 expression with the tdTomato tag as well as knockout the gene in a Cre-dependent manner.

**Supplemental Figure 2:**
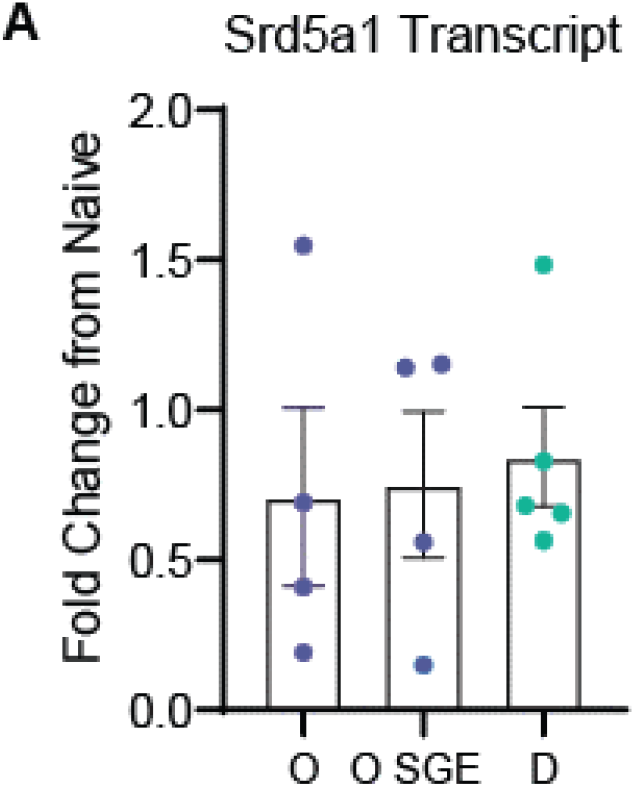
Observational fear exposure does not alter 5_α_-reductase type 1 expression. **A** Transcript levels of Srd5a1 in the BLA in observer (O), observers treated with SGE-516 (O SGE), and demonstrator (D) mice after undergoing observational fear. Transcript levels normalized to naïve average. No significant decrease from naïve or between experimental groups was found. n = 4-5 mice per experimental group. [F(2,10)=0.09150].

## Notes

### Competing Interest Statement

J.M. serves on the Scientific Advisory Board for SAGE Therapeutics and has a Sponsored Research Agreement with SAGE Therapeutics.
M.L. is employed by SAGE Therapeutics

